# DeepANIS: Predicting antibody paratope from concatenated CDR sequences by integrating bidirectional long-short-term memory and transformer neural networks

**DOI:** 10.1101/2021.08.16.456569

**Authors:** Pan Zhang, Shuangjia Zheng, Jianwen Chen, Yaoqi Zhou, Yuedong Yang

## Abstract

**Motivation:** Antibodies are a type of important biomolecules in the humoral immunity system, which can bind tightly to potential antigens with high affinity and specificity. An accurate identification of the paratope, the binding sites with antigens, is crucial for antibody mechanistic research and design. Although many methods have been developed for paratope prediction, further improvement of their accuracy is necessary.

**Results:** In this study, we concatenated the sequences of Complementarity Determining Regions (CDRs) within a single antibody to better capture nonlocal interactions between different CDRs and loop type-specific features for improving paratope prediction. We further integrated BiLSTM and transformer networks to gain the dependencies among the residues within the concatenated CDR sequences and to increase the interpretability of the model. The new method called DeepANIS (Antibody Interacting Site prediction) outperforms other antibody paratope prediction methods compared.

**Availability:** The DeepANIS method is freely available as a webserver at https://biomed.nscc-gz.cn:9094/apps/DeepANIS and for download at https://github.com/HideInDust/DeepANIS

**Supplementary information:** Supplementary data are available at *Bioinformatics* online.

## 1 Introduction

Antibodies, one of the most important biomolecules in the humoral immunity system, are responsible for neutralizing undesirable foreign molecules from pathogens such as bacteria and viruses (Frank, 2002). Such neutralization is achieved through their tight and specific binding with antigens followed by subsequent destruction mediated by the immune system. Typical antibodies are Y-shaped proteins consisting of two pairs of heavy and light chains linked by disulfide bridges. These chains are made of constant domains (C domains) that determine the functional properties, and variable domains (V domains) that are responsible for antigen binding (Moser and Leo, 2010). Each variable domain is composed of three hypervariable regions known as Complementarity Determining Regions (CDRs) and four relatively constant Framework Regions (FRs) (Schroeder Jr and Cavacini, 2010). The whole antibody contains six CDRs located within its binding loops, among which three are on the heavy chain (H1, H2, H3) and three are on the light chain (L1, L2, L3). CDRs typically contained antigen-binding residues (paratope).

Currently, the biotechnology and biopharmaceutical industries increasingly take advantage of the binding malleability of antibodies because the variability of CDR sequences enables antibodies to form complexes with almost any antigen (Ecker, et al., 2015; Reichert, 2017). With antibodies being the most promising class of biopharmaceuticals, their optimizations are often required for specific properties such as binding affinity, expression levels, stability, and solubility (Chiu and Gilliland, 2016). Such functional optimization requires the knowledge of antibody paratopes. Focusing on paratopes is important because less than one half of CDRs are involved in antigen binding (Esmaielbeiki, et al., 2016). Accurate identification of a paratope allows the separation of the binding from non-binding residues, so that further optimization can be better focused on the region of interest. Although experimental techniques can provide the gold standard for binding mode identification, they are time-consuming and labour-intensive. Thus, it is necessary to make computational prediction of paratope to complement experimental studies. Early studies on paratope prediction are mainly based on hand-coded physical models, requiring vast amounts of expert experience (Duhovny, et al., 2002; Krawczyk, et al., 2013). These methods require the structural and sequence information of antibody and antigen, limiting their usage in the real-world application of paratope identification.

More recently, deep learning techniques have been highly successful in many research areas where other traditional approaches appeared to have reached their limits. Examples are proteins structure prediction (Senior, et al., 2020), virtual screening (Zheng, et al., 2020), and antibody paratope prediction (Deac, et al., 2019; Liberis, et al., 2018). One advantage of deep learning is that it can perform automatic feature extraction directly from the original input data, thereby eliminating the need for domain experts to manually design features (Goodfellow, et al., 2016). In a previous study, the bidirectional long short-term memory neural networks (BiLSTM) (Hochreiter and Schmidhuber, 1997), which is an improvement of recurrent neural networks (RNNs) (Schuster and Paliwal, 1997), showed state-of-the-art results in many difficult sequencial problems. It can better capture the dependencies in sequences without standard directions. The transformer architecture (Vaswani, et al., 2017) further removes traditional recurrent units with significant advantages in machine translation. It employs the self-attention mechanism to extract both local and global features of sentences and increase the interpretability.

Here, we introduced a deep learning-based method for making sequence-based prediction of ANtibody Interacting Site (DeepANIS). Because it is difficult and expensive to obtain experimental structure of antibodies, only sequences and incurred properties are available in the early stages of development such as antibody discovery and antibody design, As a result, a sequence-based method is more useful in real-world applications. Unlike previous methods, we utilized CDR sequences only and concatenate those CDRs within a single antibody. Such concatenation allows our model to capture the interaction between different CDRs and loop type-specific features. In addition, the architecture of DeepANIS builds upon the bidirectional LSTM and transformer encoder, which can gain the dependencies among arbitrary residues within a concatenated CDR sequence and increase the interpretability of our model. DeepANIS was shown to outperform Parapred and ProABC (Liberis, et al., 2018; Olimpieri, et al., 2013). Parapred is a deep-learning method using convolutional and recurrent neural networks with CDRs inputted separately to predict their antibody paratope. ProABC is a random-forest-based method that employs the whole antibody sequence (no need for 3D structure) and extra features such as antigen volume and germline family.

## 2 Methods

### 2.1 Datasets

To train and test our models, we utilized a subset obtained from the Structural Antibody Database (SAbDab) (Dunbar, et al., 2014), which contains antibody and antigen crystal structures, to train and test our models. The subset was selected according to the same rules as Parapred (Liberis et al., 2018): (1) Antibodies should have variable domains of the heavy (VH) and light (VL) chains; (2) The structure resolution is better than 3Å; (3) No two antibody sequences have >95% sequence identity; (4) Each antibody has at least five residues in contact with the antigen bound to it. The final dataset contains 277 antibody-antigen bound complexes. Residues with missing electron density in the antibody sequence were assumed to be non-binding.

### 2.2 Data preprocessing and feature encoding

Our sequence-based model addresses a paratope classification task. The paratope is contained within the CDRs of the antibody or two extra residues at both ends of CDRs (Krawczyk, et al., 2013; Kunik, et al., 2012). To construct the input, we identify the CDR sequence of each antibody and the additional four extra residues using the Chothia encoding format (Al-Lazikani, et al., 1997). These extended CDR sequences are encoded as vectors prior to being processed by our model. We use an alphabet named 𝒟 to denote 21 types of residues, which contains 20 letters for 20 canonical residues and a letter ‘ X’ for non-standard residues. Then a CDR sequence can be defined as a sequence of residues *S*_*cdr =*_ (*p*_1_, *p*_2_, …, *p*_*n*_) where *p*_*i*_ ∈ 𝒟 stands for the residue at position i of the CDR sequence, and *n* represents the length of the CDR sequence (CDR sequences usually have different lengths). *S*_*cdr*_ is encoded as follows:

#### Embedding

We didn’t use one-hot encoding of residues since the sparse encoding of residues cannot be processed by a transformer encoder to calculate the attention matrix. Instead, we encoded each residue type into an integer (1-21) for its residue type and input the sequence vector to the embedding network (see below the Embedding Layer part) for embedding representations.

#### Evolutionary information

We employed the PSSM (position-specific scoring matrix) and the HMM matrix. Specifically, the HMM profile was generated by HHblits v3.0.3 in aligning the UniClust30 profile HMM database with default parameters (Mirdita, et al., 2017). PSSM was generated by PSI-BLAST v2.7.1 (Altschul, et al., 1997) using the UniRef90 sequence database after three iterations. Each residue contains 50 features with 20 from PSSM and 30 from HMM.

#### Predicted Structural properties

We obtained the predicted one-dimensional structural features from SPIDER3 (Heffernan, et al., 2017), which is one of the most accurate predictors. The feature group consists of 14 features: (1) three probability values respectively for three secondary structure states, (2) Accessible Surface Area (ASA), (3) eight values for the sine/cosine values of backbone torsion angles (phi, psi, theta, tau), (4) two values for Half-Sphere Exposures based on the C_α atom (HSE-up and HSE-down).

### 2.3 CDR concatenating

There are six CDRs located within binding loops of an antibody, three on the heavy chain (H1,H2,H3) and three on the light chain (L1,L2,L3). Different from previous studies where CDRs were individually processed (Liberis, et al., 2018), we concatenate 6 CDRs from one antibody into a sequence using five tags to capture the interaction between different CDRs (Fig S1). More specifically, for each antibody, we obtain a sequence *𝒞* = (*H*_1_*UH*_2_*UH*_3_*UL*_1_*UL*_2_*UL*_3_), in which *H*_1−3_, *L*_1−3_ denotes the six CDR sequences from different chains (heavy chain and light chain) and the letter ‘ U’ denotes the tag between different CDR sequences. Finally, we obtain protein sequences from 277 antibodies with different lengths, and padded them to the longest length in batches to speed up the training process. It is worth noting that the padding is not necessary during evaluation and prediction.

### 2.4 Model architecture

Our sequence-based neural network model consists of three components (Fig. 1). The first component is an embedding layer, which obtains trainable embedding representations of the input concatenated CDR sequences. The second component is the bidirectional LSTM and transformer encoder, which can capture the dependency between the residues within a concatenated CDR sequence. Finally, the output vector of the second component employs full connection layers to yield the prediction results of this node classification task.

**Fig. 1.**
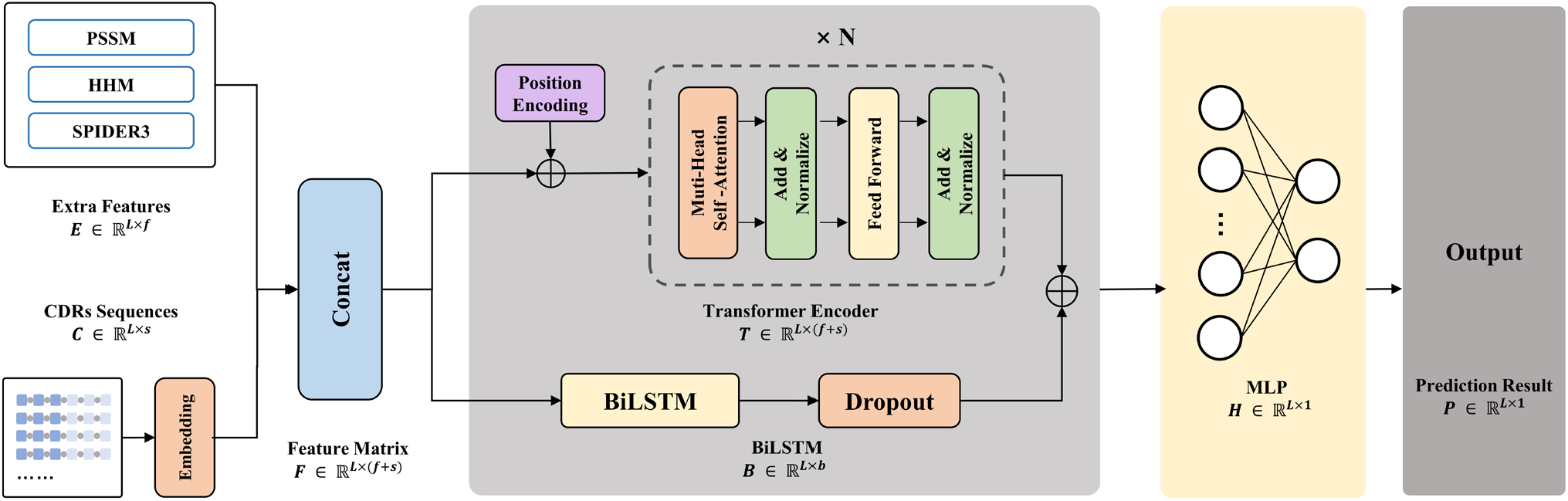
The architecture of our neural network model. For each sample, the feature matrix (F) consists of embedding features (C) and other additional features (E) is processed by BiLSTM and transformer encoder. Then the output of the second component (T) and (B) employs the full connection layer to make the final prediction.

#### 2.4.1 Embedding Layer

For each concatenated CDR sequence *𝒞* = (*H*_1_*UH*_2_*UH*_3_*UL*_1_*UL*_2_*UL*_3_), we generate a L×1 vector *𝒱* in which each element is an integer representing the residue type. The embedding layer initialize a 22×*d* vector *W*, which contains 22 rows (0 for padding and 1-21 for residue type). Then embedding layer process *𝒱* to a L×22 vector *𝒱*′, in which each row is one hot encoding of the corresponding residue denoted by an integer. At last, the embedding layer computes trainable embedding representations of *𝒞*, i.e.,

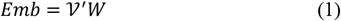

### 2.4.2 Bidirectional LSTM

We utilize the Long Short-Term Memory (LSTM) to learn long-range dependencies in concatenated CDR sequences. LSTM cell is a special RNN (recurrent neural network) cell, which consists of the following computation steps:

The first step in the LSTM cell is to determine what information needs to be discarded, which is implemented with a forget gate. The forget gate looks at *h*_*t*−1_ (previous output) and *x*_*t*_ (input of current time step), and uses a sigmoid function *σ* to output a 0-1 number *f*_*t*_, i.e.,

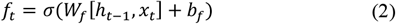

where *W* and *b* are trainable parameters of our model, [*h*_*t*−1_, *x*_*t*_] means to concatenate *h*_*t*−1_ and *x*_*t*_.

The next step in the LSTM cell is to determine what information needs to be stored, which is implemented with an input gate. The input gate uses a sigmoid function *σ* to decide what values we will update (*i*_*t*_) and a function *tanh* to create a candidate value *j*_*t*_ that will be added to the state, i.e.,

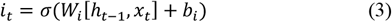

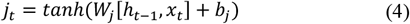

Then the LSTM cell updates the old cell state *C*_*t*−1_ into the new cell state *C*_*t*_ using *f*_*t*_, *i*_*t*_, *j*_*t*_, i.e.,

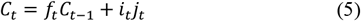

Finally, the LSTM cell uses a sigmoid function to decide what part of *C*_*t*_ will be output (*O*_*t*_), then it puts *C*_*t*_ through function *tanh* weighted by *O*_*t*_ to calculate the final output *h*_*t*_, i.e,

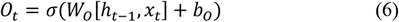

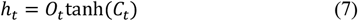

Because the concatenated CDR sequences do not have a canonical direction, a bidirectional LSTM (BiLSTM) is utilized to process them:

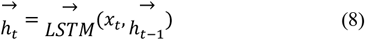

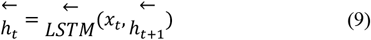

where 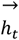 and 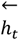 are concatenated as the output of time step t, which is a more information-enriched vector than the input *x*_*t*_:

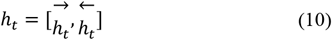

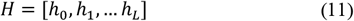

We denote the output of BiLSTM layer as H for simplicity. If the hidden unit number of each BiLSTM cell is *u*, the shape of *H* will be *L*-by-2*u*.

### 2.4.3 Transformer Encoder

To gain more dependency between the residues within a concatenated CDR sequence and increase the interpretability of our model, we apply a transformer encoder in our model. The architecture of the transformer encoder is shown in Fig.2. Several identical layers are stacked in the encoding phase. Each layer is composed of a multi-head self-attention sublayer and a feedforward network (FFN) sublayer. The two sub-layers are integrated using the residual connection and layer normalization.

**Fig. 2.**
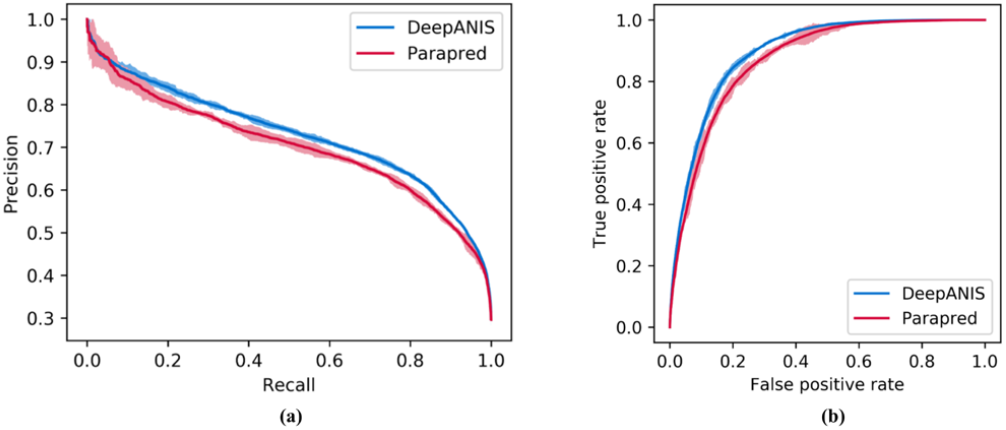
The receiver operating characteristics curves (a) and precision-recall curves (b) given by Parapred and DeepANIS, respectively. Error bars show 95% confidence bounds.

A multi-head self-attention unit contains several attention layers that can execute the attention mechanism in parallel, and then concatenate these layers and project them to the final values. The input of the scaled-dot attention layers are three matrices: the query (Q), the key (K), and the value (V), which are created by multiplying the input feature matrix *F* by three trainable weight matrices during the training process. We then calculate the attention weight for each residue within a concatenated CDR sequence as follows:

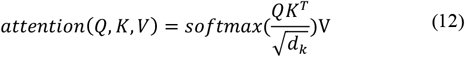

where *d*_*k*_ is a scaling factor, depending on the size of feature matrix and head. *QK*^*T*^ computes the degree of association between keys and queries. The large dot product means that keys and queries are aligned well. Then we obtained an attention map, using the *softmax* function. Finally, we multiplied the attention map by the value vector. Using this procedure, the transformer encoder important features from the source concatenated CDR sequences.

The transformer encoder lacks a way to explain the order of the residues within an input sequence because it removes the recurrent units. To solve this, we employed the position encoding proposed in the previous study (Vaswani, et al., 2017):

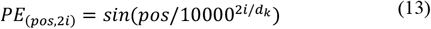

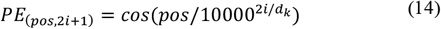

where *pos* is the position and *i* is the dimensional index of position encoding matrix (the same dimension as the input feature matrix).

#### 2.4.4 Multilayer Perceptron

The output of the transformer encoder and BiLSTM are concatenated and then is fed to the multilayer perceptron (MLP) to obtain final prediction results:

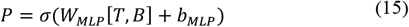

where *T* is the output of transformer encoder, *B* is the output of BiLSTM, *W*_*MLP*_ and *b*_*MLP*_ are trainable parameters of MLP, *σ* is the sigmoid function that maps the value in *P* to 0-1 for prediction.

### 2.5 Training and Evaluation

#### 2.5.1 Loss Functions

We used the backpropagation algorithm (Rumelhart, et al., 1985) and the gradient descent method to train all parameters of our model, and the loss function is the binary cross-entropy function (De Boer, et al., 2005):

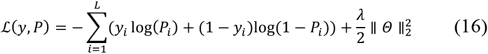

where *y*_*i*_ and *p*_*i*_ are true and predicted labels of the *i*th residue in the concatenated CDR sequences, respectively, *Θ* is the set of all weight and bias parameters in our model, *λ* is the L2 regularization hyperparameter. The adam optimizer were used in this study (Kingma and Ba, 2014). The DeepANIS has been implemented using Keras library 2.2.4 and Tensor-flow-GPU 1.9.0 (Abadi, et al., 2016). All the training and testing processes were performed on GTX 1080Ti GPU.

#### 2.5.2 Hyper-parameter-tuning

Our model includes multiple hyperparameters. We performed a hyperparameter search to obtain the best parameters:

1. **Hidden unit number of BiLSTM:** The dimension of hidden states depends on the hidden unit number of BiLSTM. We varied the hidden unit number of BiLSTM {16,32,64,128,256,512} and found that 256 units provided the best performance.
2. **Transformer encoder layers:** Several identical layers of a transformer encoder are stacked in the encoding phase. A high number of transformer encoder layers means the deeper and wider information mined from concatenated CDR sequences. However, excessive layers will cause greater algorithm complexity and a performance reduction. Therefore, it is crucial to choose a balanced number of layers. We varied the number of transformer encoder layers from 1 to 6 and found that 2 layers provided the best performance.
3. **Attention heads:** The attention map learned by the transformer encoder provides weight coefficients, which can focus on key residues of the concatenated CDR sequences. Different attention heads enable the attention of input sequences from different views. We varied the number of attention heads {1,2,4,8} and found that 4 attention heads provided the best performance.

#### 2.5.3 The 10-fold cross-validation

To ensure an unbiased evaluation and obtain statistically significant results, we use the 10-fold cross-validation technique on our dataset (277 complexes). This technique divides the dataset into 10 subsets, 9 subsets of which was employed as the training set and the remaining portion as the test set. The test subset is rotated so that all structures were employed in the test. Each test will yield binding probabilities, the combined results are used as an estimate of the algorithm performance. Ten folds were randomly divided 10 times to examine the robustness of the training.

#### 2.5.4 Performance measure

Our model was evaluated by the following indicators: MCC (Matthews Correlation Coefficient), AUPR (the area under the precision-recall curve), AUROC (the area under the receiver-operating characteristic) (Lobo, et al., 2008), and F1-score, which are defined as:

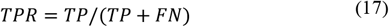

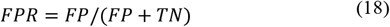

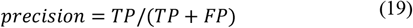

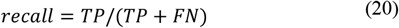

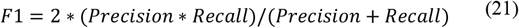

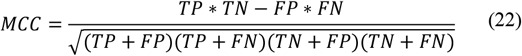

where *TP, FP, FN*, and *TN* denote the number of true positives, false positives, true negatives and false negatives, respectively, *FPR* and *TPR* are the x and y axis of the receiver-operating characteristic, *precision* and *recall* are the x and y axes of the precision recall curve.

## 3 Results and Discussion

### 3.1 Performances on the 10-fold cross-validation

The results of ten fold cross validations are shown in Table.1. DeepANIS achieved a MCC score of 0.606 ± 0.002, and an AUPR score of 0.727±0.004 in the 10-fold cross-validations. In order to illustrate the importance of various features, we evaluated them individually and by addition and removal of each feature group. As shown in Table 1, sequence profiles PSSM yielded the best performance (a MCC score of 0.597±0.005, and an AUPR score of 0.724±0.003) in the 10-fold cross-validation. The performance of the other evolution-based feature (the HMM profile) is similar but slightly worse than PSSM. The predicted structural features by SPIDER3 achieved the worst performance with MCC of 0.583±0.004 and AUPR of 0.711±0.004. The removal and addition of each feature group has the similar trend: PSSM makes the largest contribution, followed by HMM and SPIDER3. Nevertheless, each feature group contributes positively to the overall performance and thus, all these features were utilized in our final method.

**Table 1.** Performance of DeepANIS in 10-fold cross-validation by different feature groups individually or in combination. To convert the predicted binding probabilities to binary labels, we used thresholds in the range of [0.482, 0.523] obtained by maximizing Youden’s index (Youden, 1950); the labels were used to compute the MCC and F-score metrics. The AUC and F-score metrics are shown in Supplementary.

| Feature groups<sup>a</sup> | MCC | AUPR | Feature groups<sup>b</sup> | MCC | AUPR | Feature groups<sup>c</sup> | MCC | AUPR |
| --- | --- | --- | --- | --- | --- | --- | --- | --- |
| (Embedding) | $0.585 \pm 0.003$ | $0.714 \pm 0.003$ | **DeepANIS** | **$0.606 \pm 0.002$** | **$0.727 \pm 0.004$** | Embedding | $0.585 \pm 0.003$ | $0.714 \pm 0.003$ |
| PSSM | **$0.597 \pm 0.005$** | **$0.724 \pm 0.003$** | -PSSM | $0.597 \pm 0.003$ | $0.723 \pm 0.002$ | +PSSM | $0.597 \pm 0.005$ | $0.724 \pm 0.003$ |
| HMM | $0.591 \pm 0.003$ | $0.716 \pm 0.006$ | -HMM | $0.601 \pm 0.003$ | $0.723 \pm 0.003$ | +HMM | $0.603 \pm 0.002$ | $0.725 \pm 0.003$ |
| SPIDER3 | $0.583 \pm 0.004$ | $0.711 \pm 0.004$ | -SPIDER3 | $0.603 \pm 0.002$ | $0.725 \pm 0.003$ | + SPIDER3 | **$0.606 \pm 0.002$** | **$0.727 \pm 0.004$** |

<sup>a</sup> Performances based on individual feature groups, <sup>b</sup> by removing each feature group from all feature groups, and <sup>c</sup> by adding feature groups recursively.

### 3.2 Ablation study

#### 3.2.1 CDR concatenation improves prediction performance

We used a neural network model (CNN+BiLSTM) to assess the effect of CDR concatenation. As shown in Table 2A, Using concatenated CDR sequences yields statistically significant improvement over inputting CDR individually. More specifically, the concatenating method achieved a MCC of 0.574 ± 0.004, an AUPR of 0.706 ± 0.005, an AUC of 0.888±0.002, and a F-score of 0.694±0.002. By the comparison, the baseline approach (CDRs processing individually method under the same neural network) leads to a MCC of 0.556±0.007, an AUPR of 0.700±0.006, an AUC of 0.878±0.003, and a F-score of 0.688±0.007. Thus, CDR concatenating was effective for improving prediction of binding sites of antibodies.

**Table 2.** The results of Ablation study. (**A)** Performance comparison between inputting concatenated CDRs and inputting individual CDRs. A CNN+BiLSTM model was used for this comparison. (**B)** Performance comparison among different neural network architectures. All models employed the same embedding features from concatenated CDR sequences as input. The Transformer+BiLSTM is the model architecture for DeepANIS.

| Method<sup>A</sup> | MCC | AUPR | AUC | F-score | Network<sup>B</sup> | MCC | AUPR | AUC | F1-score |
| --- | --- | --- | --- | --- | --- | --- | --- | --- | --- |
| Individually | 0.556±0.007 | 0.700±0.006 | 0.878±0.003 | 0.688±0.007 | BiLSTM | 0.572±0.001 | 0.705±0.004 | 0.888±0.001 | 0.692±0.002 |
| Concatenated | **0.574±0.004** | **0.706±0.005** | **0.888±0.002** | **0.694±0.002** | Transformer | 0.527±0.002 | 0.648±0.004 | 0.871±0.002 | 0.645±0.003 |
| Individually: The 6 CDRs from one antibody are inputted individually. |  |  |  |  | **Transformer**<br>**+ BiLSTM** | **0.585±0.003** | **0.714±0.003** | **0.889±0.001** | **0.701±0.002** |
| Concatenated: the 6 CDRs from one antibody are concatenated into a single sequence. |  |  |  |  |  |  |  |  |  |

#### 3.2.2 Transformer encoder improves prediction performance

To illustrate the usefulness of a transformer encoder, we made a comparative study using different neural network architectures. As shown in Table 2B, using the same embedding features from concatenated CDR sequences as input, the single transformer-encoder model can achieve a MCC of 0.527±0.002, an AUPR of 0.648±0.004, an AUC of 0.871±0.002, and a F-score of 0.645±0.003. This is slightly worse than the performance given by the single BiLSTM model. Our model (Transformer+BiLSTM) achieves the best performance with a MCC of 0.585±0.003, an AUPR of 0.714±0.003, an AUC of 0.889±0.001, and a F-score of 0.701±0.002. The result confirms the usefulness of the transformer encoder for improving prediction of antibody binding sites

### 3.3 Comparisons with other methods

Table 3 compares DeepANIS with proABC, Parapred, and the netweorks based on CNN and BiLSTM. The random-forest method proABC, achieved a MCC score of 0.522 and an AUC score of 0.851. Parapred, a CNN+BiLSTM based method, achieved a MCC score of 0.554±0.009, an AUPR score of 0.701±0.008, an AUC score of 0.878±0.004, and a F-score of 0.690±0.006. To ensure a fair comparison, we also emplolyed a network model of CNN+BiLSTM to mimic Parapred. An essentially identical result to Parapred was obtained. By comparison, DeepANIS performs the best with or without additional features besides embedding features. The improvement over Parapred by DeepANIS is further confirmed by the ROC curves and the precision-recall curves as shown in Figure 2. These curves given by DeepANIS are mostly above those given by Parapred, indicting a robust improvement at different levels of sensitivities.

**Table 3.** Performance comparison among different neural network architectures. All models employed the same embedding features from concatenated CDR sequences as input. The Transformer+BiLSTM is the model architecture for DeepANIS.

| Method | MCC | AUPR | AUC | F-score |
| --- | --- | --- | --- | --- |
| proABC | 0.522 | — | 0.851 | — |
| Parapred | $0.554 \pm 0.009$ | $0.701 \pm 0.008$ | $0.878 \pm 0.004$ | $0.690 \pm 0.006$ |
| CNN+BiLSTM | $0.556 \pm 0.007$ | $0.700 \pm 0.006$ | $0.878 \pm 0.003$ | $0.688 \pm 0.007$ |
| **DeepANIS (embedding features)** | **$0.585 \pm 0.003$** | **$0.714 \pm 0.003$** | **$0.889 \pm 0.001$** | **$0.701 \pm 0.002$** |
| **DeepANIS (all features)** | **$0.606 \pm 0.002$** | **$0.727 \pm 0.004$** | **$0.896 \pm 0.001$** | **$0.717 \pm 0.002$** |

### 3.4 Transformer encoder finds important local features in CDR sequence

Another benefit of using a transformer encoder is its interpretability, which can help us investigate what our neural network model has learned. We chose one fold from 10-fold cross-validation randomly for this analysis. For an input sequence of length is L, the transformer encoder can yield an *L* × *L* attention map *A*_*tt*_, in which *Att*_*ij*_ denotes the attention score of the *j*th residue to the *i*th residue. A higher attention score indicates a closer correlation between the *j*th amino and the *i*th residue. We calculated the sum of each column from *Att* without the diagonal elements. A larger sum of the columns for a residue indicates the more attention paid to the residue by the model. We chose top *N* residues with the highest attention scores from each test sample, and then analyzed their final prediction results compared with the BiLSTM single model (*N*=2, 4, 6, 8, 10). Figure 3 shows that the difference between DeepANIS and Parapred is smaller for residues with low attenstion scores. That is, the transformer encoder can focus its attention on some residues within concatenated CDR sequences and thereby improve prediction by integrating their features in multi-head attention units. To further illustrate this, we selected one complex structure in the test set whose pdb id is 2jel. Figure 4 shows the attention map obtained along with the complex structures with true positives, false positives, true negatives and false negatives highlighted in different colors on the CDR regions for DeepANIS and Parapred. The main difference between the two methods are that BiLSTM in Parapred yields more false positives. For example, positions 1, 6, and 7 of the second light chain (L2) and position 3 of the first heavy chain (H1) were predicted correctly as non-binding residues by DeepANIS but incorrectly as binding residues by Parapred. Examining the attention map reveals that the attention weights of these 4 residues ranked 3rd, 10th, 11th, and 13th among the 79 residues of the concatenated CDR sequence. Thus, the additional attention of the transformer encoder to the four residues leads to their correct classification as non-binding residues.

**Fig.3.**
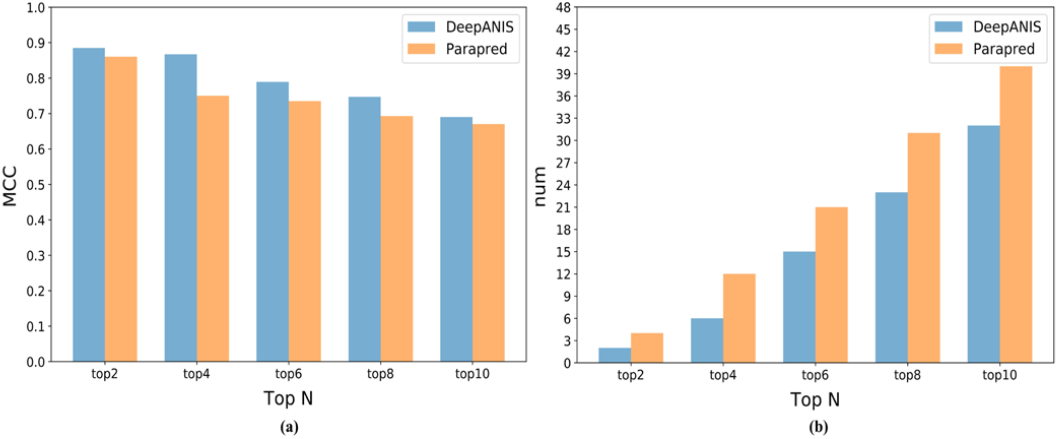
Comparison of the indicators of the top-k noted residues selected by transformer encoder. The MCC (a) and FPN (b) of our method were better than BiLSTM single model, and the difference between the two methods decreased with the decrease of attention score, which is in line with expectations.

**Fig.4.**
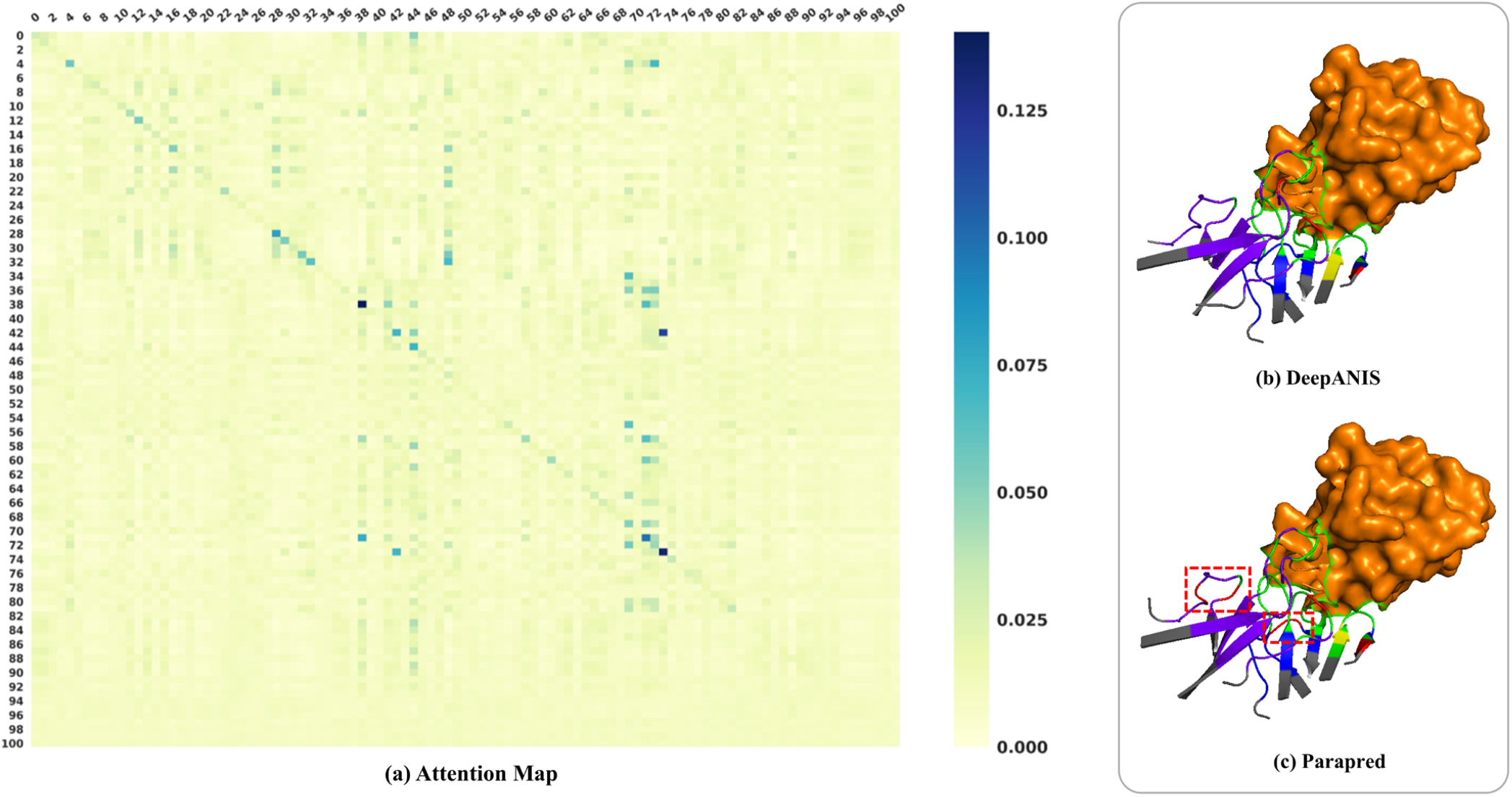
The visualization results of 2jel complex. The attention map **(a)** was obtained by transformer encoder, in which each position denoted the attention score between two residues. **(b)** and **(c)** are the visualization results of our method and LSTM method respectively. Different colors represent the following meanings: purple: true negatives of light chain; blue: true negatives of heavy chain; yellow: false negatives; green: true positives; red: false positives. The main difference between results predicted by two methods was the distribution of false positives.

## 4 Conclusions

In this work, we introduced a sequence-based method DeepANIS (ANti-body Interacting Site) dedicated to the prediction of antibody paratope. Compared to other methods, we concatenated CDRs from a single anti-body, which allows us to capture nonlocal interactions among CDRs. The combined use of the BiLSTM and the transformer encoder further permits an improved attention to specific residues for better classification. The results indicate that the attention map obtained from the transformer encoder may be useful to investigate the binding mechanisms between antibodies and antigens, which is a subject for further research.

## Acknowledgements

This study has been supported by the National Key R&D Program of China (2020YFB0204803), National Natural Science Foundation of China (61772566), Guangdong Key Field R&D Plan (2019B020228001 and 2018B010109006), Introducing Innovative and Entrepreneurial Teams (2016ZT06D211), Guangzhou S&T Research Plan (202007030010). The support of Shenzhen Science and Technology Program (Grant No. KQTD20170330155106581) and the Major Program of Shenzhen Bay Laboratory S201101001 is also acknowledged. This work is also supported in part by the supercomputing facility of the Shenzhen Bay Laboratory.

