## Supplementary material for "DeepANIS: Predicting antibody paratope from concatenated CDR sequences by integrating bidirectional long-short-term memory and transformer neural networks": (Fig S1)

**Table S1.** The abbreviations listed in this paper.

| Abbreviation | Full Name |
|---|---|
| AUPR | Area under the precision-recall curve |
| AUROC | Area under the receiver operating characteristic curve |
| LSTM | Long short-term memory |
| CDR | Complementarity determining regions |
| CNN | Convolutional neural network |
| CV | Cross-validation |
| HMM | Hidden Markov models |
| BiLSTM | Bidirectional long short-term memory |
| MCC | Matthews correlation coefficient |
| MLP | Multilayer perceptron |
| PDB | Protein Data Bank |
| PSSM | Position-specific scoring matrix |
| RF | Random forest |
| RNN | Recurrent neural network |
| SAbDab | Structural antibody database |

**Table S2.** The performance of DeepANIS on CV using different hyperparameters.

| Hyperparameter | Value | MCC | AUPR | AUC | F-score |
|---|---|---|---|---|---|
| Hidden units of BiLSTM | 16 | 0.585 | 0.71 | 0.875 | 0.695 |
| | 32 | 0.592 | 0.72 | 0.885 | 0.708 |
| | 64 | 0.596 | 0.723 | 0.892 | 0.714 |
| | 128 | 0.6 | 0.725 | 0.89 | 0.712 |
| | **256** | 0.605 | 0.727 | 0.897 | 0.718 |
| | 512 | 0.602 | 0.728 | 0.895 | 0.716 |
| Transformer encoder layers | 1 | 0.601 | 0.722 | 0.892 | 0.711 |
| | **2** | 0.606 | 0.726 | 0.898 | 0.716 |
| | 3 | 0.604 | 0.726 | 0.895 | 0.715 |
| | 4 | 0.595 | 0.717 | 0.891 | 0.709 |
| | 5 | 0.591 | 0.712 | 0.886 | 0.704 |
| | 6 | 0.583 | 0.706 | 0.872 | 0.696 |
| Attention heads | 1 | 0.586 | 0.708 | 0.871 | 0.694 |
| | 2 | 0.598 | 0.719 | 0.892 | 0.711 |
| | **4** | 0.604 | 0.725 | 0.895 | 0.717 |
| | 8 | 0.603 | 0.721 | 0.891 | 0.712 |

We concatenate 6 CDRs from one antibody into a sequence using five tags to capture the interaction between different CDRs. An example of concatenated CDR sequences is shown in Figure S1. The $H_{1-3}$ and $L_{1-3}$ denotes the six CDR sequences from different chains of a single antibody. The tags denoted by 'U' aim to connect and distinguish different CDR sequences. Each amino acid residue within the CDR sequences is encoded to a feature matrix, which consists of embedding and other additional features. Then, each concatenated CDR sequence is padded to the length of the longest sequence for training in batches. The final dataset contains 277 concatenated CDR sequences.

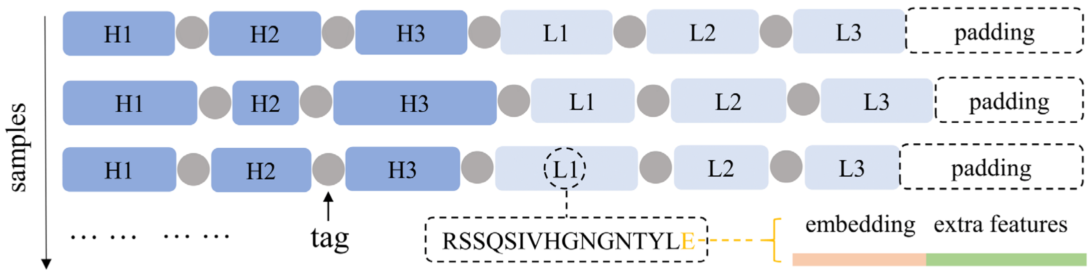


**Figure S1.** An example of concatenated CDR sequences.
